# Cellular heterogeneity during mouse pancreatic ductal adenocarcinoma progression at single-cell resolution

**DOI:** 10.1101/539874

**Authors:** Abdel Nasser Hosein, Huocong Huang, Zhaoning Wang, Kamalpreet Parmar, Wenting Du, Jonathan Huang, Anirban Maitra, Eric Olson, Udit Verma, Rolf A. Brekken

## Abstract

**Background & Aims:** Pancreatic ductal adenocarcinoma (PDA) is a major cause of cancer-related death with limited therapeutic options available. This highlights the need for improved understanding of the biology of PDA progression. The progression of PDA is a highly complex and dynamic process featuring changes in cancer cells and stromal cells; however, a comprehensive characterization of PDA cancer cell and stromal cell heterogeneity during disease progression is lacking. In this study, we aimed to profile cell populations and understand their phenotypic changes during PDA progression.

**Methods:** We employed single-cell RNA sequencing technology to agnostically profile cell heterogeneity during different stages of PDA progression in genetically engineered mouse models.

**Results:** Our data indicate that an epithelial-to-mesenchymal transition of cancer cells accompanies tumor progression. We also found distinct populations of macrophages with increasing inflammatory features during PDA progression. In addition, we noted the existence of three distinct molecular subtypes of fibroblasts in the normal mouse pancreas, which ultimately gave rise to two distinct populations of fibroblasts in advanced PDA, supporting recent reports on intratumoral fibroblast heterogeneity. Our data also suggest that cancer cells and fibroblasts are dynamically regulated by epigenetic mechanisms.

**Conclusion:** This study systematically outlines the landscape of cellular heterogeneity during the progression of PDA. It strongly improves our understanding of the PDA biology and has the potential to aid in the development of therapeutic strategies against specific cell populations of the disease.

## Introduction

Pancreatic ductal adenocarcinoma (PDA) carries the highest mortality rate of all major malignancies in industrialized countries, with a 5-year survival of 8.5%. Patients are faced with limited treatment options that achieve poor durable response rates, highlighting the need for an improved understanding of PDA disease biology [1]. PDA progression is a complex and dynamic process that requires interaction between cancer cells and stromal cells [2]. It is characterized by the formation of a unique microenvironment consisting of heterogeneous stromal cell populations that include fibroblasts, macrophages, lymphocytes, and endothelial cells. These stromal compartments are critical in driving PDA biology [3].

The dynamic phenotypic changes in different cell populations during PDA progression is not fully understood. Gene expression profiling of bulk tissues provides a limited picture of the cellular complexity of the heterogeneous cell populations in PDA. In contrast, single-cell RNA sequencing (scRNA-seq) has the potential to enable gene expression profiling at the level of the individual cell [4] and provides a powerful tool to understand the cellular heterogeneity of PDA. We applied scRNA-seq to investigate gene expression changes of cancer cells and stromal cells during PDA progression in genetically engineered mouse models (GEMMs). This unbiased approach provided evidence of considerable intratumoral cellular heterogeneity, including molecular insights into epithelial and mesenchymal populations of cancer cells and distinct molecular subtypes of macrophages and cancer-associated fibroblasts (CAFs).

## Methods

### Animal studies

*KIC, KPC* and *KPfC* mice were generated as previously described [5–7]. Mice were sacrificed when they were moribund: 60 days old for the *KIC* (n = 3, late PDA) and *KPfC* (n = 1) or 6 months old for the *KPC* (n = 1). The 2 *KIC* mice were sacrificed at 40 days old (early PDA) and “normal pancreas” mice (n = 2) were sacrificed at 60 days old. In experiments using more than one mouse, tissues were pooled prior to enzymatic digestion. The *KPfC* mouse had a pure C57BL/6 genetic background and all others had a mixed background (C57BL/6 with FVB). Ultrasound imaging was carried out under general anesthesia with isoflurane. Mice were euthanized by cervical dislocation under anesthesia. AVMA Guidelines for the Euthanasia of Animals were strictly followed. Tissues were either fixed in 10% formalin for immunohistochemistry or enzymatically digested for single-cell analysis.

### Tissue digestion

A 10x digestion buffer was prepared in PBS: collagenase type I (450 units/ml, Worthington Biochemical, Lakewood, NJ), collagenase type II (150 units/ml, Worthington), collagenase type III (450 units/ml, Worthington), collagenase type IV (450 units/ml, Gibco/Thermo Fisher, Waltham, MA), elastase (0.8 units/ml, Worthington), hyaluronidase (300 units/ml, Sigma-Aldrich, St. Louis, MO), and DNase type I (250 units/ml, Sigma-Aldrich). Tumors and pancreas were enzymatically digested into a single-cell suspension. Briefly, freshly dissected tissue was placed into a 10-cm tissue culture dish and a sterile razor blade was used to cut the tissue into fine pieces. Samples were resuspended in PBS and washed twice by centrifuge at 2000 rpm for 3 minutes and added to a 50 ml tube containing 1x digestion buffer containing 1% FBS. The tube was incubated on a shaker at 37°C for 60 minutes. Then 35 ml of PBS was added and cells were washed three times prior to filtering out debris using a 70 μm mesh filter. Single cells were resuspended in 100 μl of PBS in preparation for single-cell library creation. Cell viability was measured by trypan blue. Viability was 80% for the normal pancreas and late *KIC* samples, 75% for the early *KIC* and *KPfC*, and 90% for the *KPC*.

### Single-cell cDNA library preparation and sequencing

Library generation was performed using the 10x Chromium System (10X Genomics Inc., Pleasanton, CA). Single-cell suspensions were washed in 1x PBS (calcium- and magnesium-free) containing 0.04% weight/volume bovine serum albumin (400 μg/ml) and brought to a concentration of 200-700 cells/μl. The appropriate volume of cells was loaded with Single Cell 3’ gel beads into a Single Cell A Chip and run on the Chromium Controller. Gel bead in emulsion (GEM) was incubated and then broken. Silane magnetic beads were used to clean up the GEM reaction mixture. Read 1 primer sequence was added during incubation and full-length, barcoded cDNA was amplified by PCR after cleanup. Sample size was checked on an Agilent Tapestation 4200 (Agilent, Santa Clara, CA) using DNAHS 5000 tape and concentration determined by a Qubit 4 Fluorometer (Thermo Fisher) using the DNA HS assay. Samples were enzymatically fragmented and underwent size selection before proceeding to library construction. During library preparation, Read 2 primer sequence, sample index, and both Illumina adapter sequences were added. Samples were cleaned up using AMPure XP beads (Beckman Coulter, Brea, CA) and post-library preparation quality control was performed using DNA 1000 tape on the Agilent Tapestation 4200. The final concentration was ascertained using the Qubit 4 Fluorometer DNA HS assay. Samples were loaded at 1.5 pM and run on the Illumina NextSeq500 High Output Flowcell (Illumina, San Diego, CA) using V2.5 chemistry. The run configuration was 26 x 98 x 8.

### Bioinformatic analyses

We used Cell Ranger version 1.3.1 (10x Genomics) to process raw sequencing data and the R-package Seurat version 2.0 [8] for downstream analyses. Cell clusters were identified via the FindClusters function using a resolution of 0.6 for all samples, using a graph-based clustering algorithm implemented in Seurat. Marker genes for each cluster were computed, and expression levels of several known marker genes were examined. Different clusters expressing known marker genes for a given cell type were selected and combined as one for each cell type. Gene ontology and pathway analysis were performed using the DAVID bioinformatics suite, version 6.8 [9].

### Histological analysis

Formalin-fixed tissues were embedded in paraffin and cut in 5 μm sections. Sections were evaluated by H&E and immunohistochemical analysis using antibodies specific for vimentin (5741, Cell Signaling Technology, Danvers, MA), BRD4 (AB128874, Abcam, Cambridge, MA), Sox9 (AB5535, EMD Millipore, Burlington, MA), CDH11 (NBP2-15661, Novus Biologicals, Centennial, CO), and H3K27ac (AB4729, Abcam). Following an initial antigen retrieval with Tris-EDTA-glycerol (10%) buffer and inhibition of endogenous peroxidase activity, the slides were incubated with primary antibody overnight at 4°C. Slides were then incubated with horseradish peroxidase or alkaline phosphatase conjugated secondary antibody (Vector Laboratories, Beringame, CA) for 1 hour at 25°C. This was followed by development using the appropriate chromogenic substrate: DAB, Warp Red or Ferangi Blue (Biocare Medical, Pacheco, CA). In the case of multichannel immunohistochemistry, slides were subsequently stripped using a sodium citrate buffer and by boiling at 110°C for 3 minutes. The procedure was then repeated as above using a different-colored chromogen for development. All human PDA samples were provided by the UT Southwestern Tissue Management Shared Resource and their use was approved by the UT Southwestern institutional review board for the purpose of research. All patient samples were de-identified and interpreted by a board-certified pathologist (KP).

## Results

### Cellular heterogeneity during PDA progression

We sought to determine the composition of single pancreatic cancer cells during progression in GEMMs. Normal mouse pancreas, 40-day-old *KIC* (*Kras^LSL-G12D^*; *Cdkn2a^flox/flox^*; *Ptf1a*^*Cre*/+^) mouse pancreas, termed “early *KIC*” (with early neoplastic changes confirmed by ultrasound; Supplementary Fig. 1), and 60-day-old *KIC* pancreas, termed “late *KIC*” (Fig. 1A) were freshly isolated and enzymatically digested followed by single-cell cDNA library generation using the 10x Genomics platform [10]. Libraries were subsequently sequenced at a depth of more than 10^5^ reads per cell. We performed stringent filtering, normalization, and graph-based clustering, which identified distinct cell populations in the normal pancreas and each stage of PDA.

**Figure 1.**
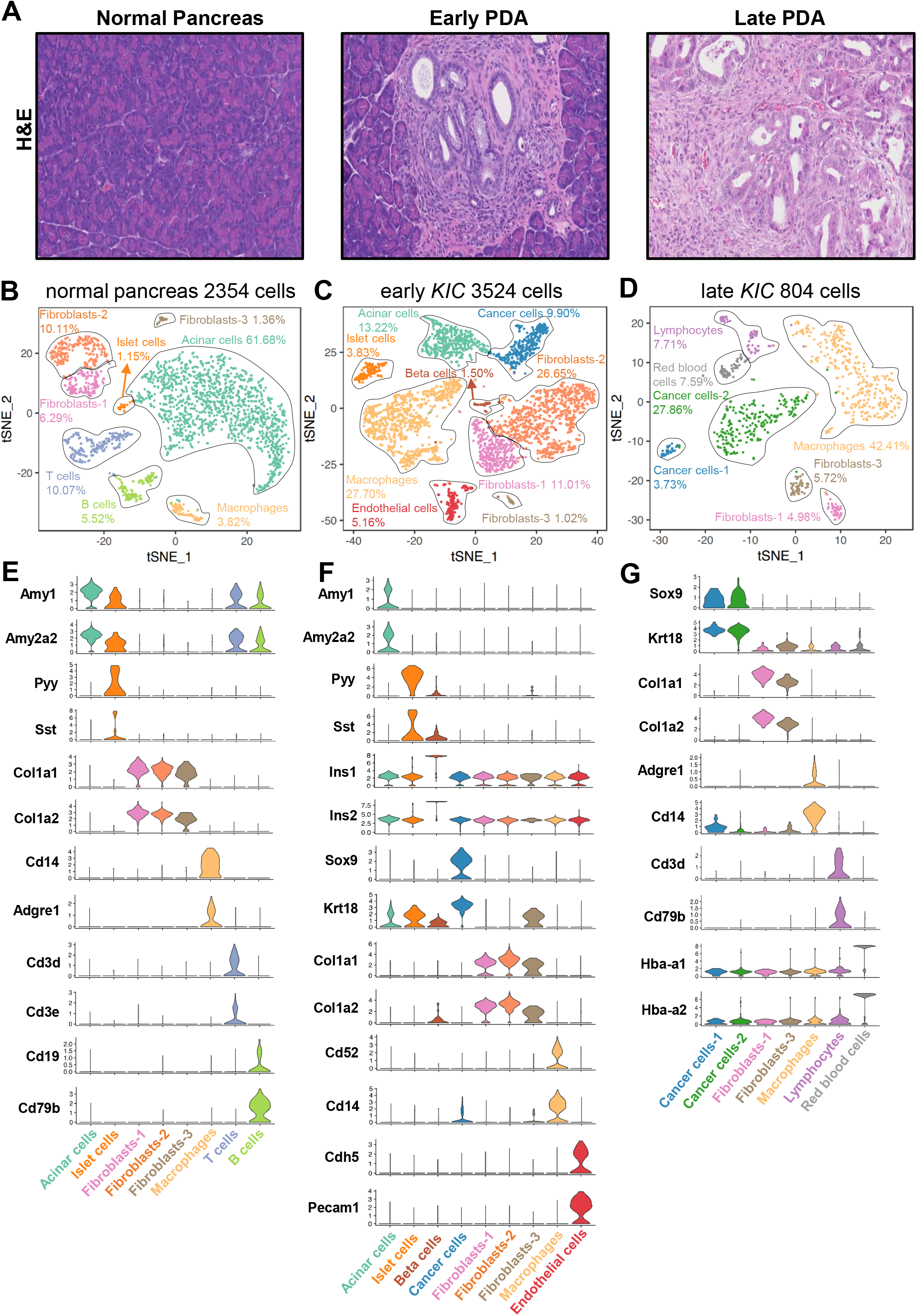
Cellular heterogeneity during PDA progression. A) Representative H&E sections of the normal pancreas, early *KIC* lesion, and late *KIC* lesion (magnification: 20x). B) tSNE plot of normal pancreas displaying 2354 cells comprising 8 distinct cell populations. C) tSNE plot of the early *KIC* lesion displaying 3524 cells containing 9 cells types with the emergence of the cancer cell population. D) tSNE plot of the late *KIC* tumor showing 804 cells and 7 distinct populations. Stacked violin plots of representative marker gene expression for each of the cell populations seen in the E) normal pancreas, F) early *KIC* lesions, and G) late *KIC* lesions.

In the normal mouse pancreas, 2354 cells were sequenced and classified into appropriate cell types based on the gene expression of known markers: acinar cells, islet cells, macrophages, T cells, and B cells, as well as three distinct populations of fibroblasts. Fibroblasts-1, fibroblasts-2, and fibroblasts-3 (Fig. 1B and E) were noted. In the early *KIC* pancreas (3524 cells sequenced), the emergence of a cancer cell population was observed (9.9% of cells), expressing known PDA markers such as *Krt18* and *Sox9* [11] (Fig. 1C and F). The acinar cell population was substantially reduced, while there was a marked increase in total macrophages and fibroblasts. Of note, the same three populations of fibroblasts seen in the normal pancreas were identified in the early *KIC* lesion. Additionally, endothelial cells were observed at this stage. This indicates that the expansion of fibroblasts and macrophages is an early event during PDA development, accompanying tumor initiation. We next characterized the late *KIC* pancreas (804 cells sequenced) and noted the absence of normal exocrine (acinar) and endocrine (islet) cells (Fig. 1D and G). Instead, two distinct populations of cancer cells were present, suggesting phenotypic cancer cell heterogeneity as a late event in the course of the disease. We also observed the presence of only two distinct fibroblast populations, which had a similar percentage in relation to total cells. Noticeably, macrophages became a predominant cell population in the late *KIC* tumor. Moreover, we observed lymphocytes at this stage. The cellular heterogeneity in cancer cells and stromal cells in the early and late *KIC* lesions highlighted the dynamic cellular changes that occur during PDA progression.

### Mesenchymal cancer cells emerge in advanced PDA

Gene expression analysis of cancer cell epithelial markers *(Cdh1, Epcam, Gjb1, and Cldn3)* and mesenchymal markers (*Cdh2, Cd44, Axl, Vim, and S100a4*) revealed that early *KIC* cancer cell populations assumed an epithelial expression profile (Fig. 2A and C). This is in contrast to tumor cell populations in the late *KIC* tumors, where we identified two distinct cancer cell populations: one enriched for epithelial markers and the other, more abundant population, enriched for mesenchymal markers (Fig. 2B and C). These data support that tumor cell epithelial plasticity contributes to cancer cell heterogeneity during the progression of *KIC* tumors.

**Figure 2.**
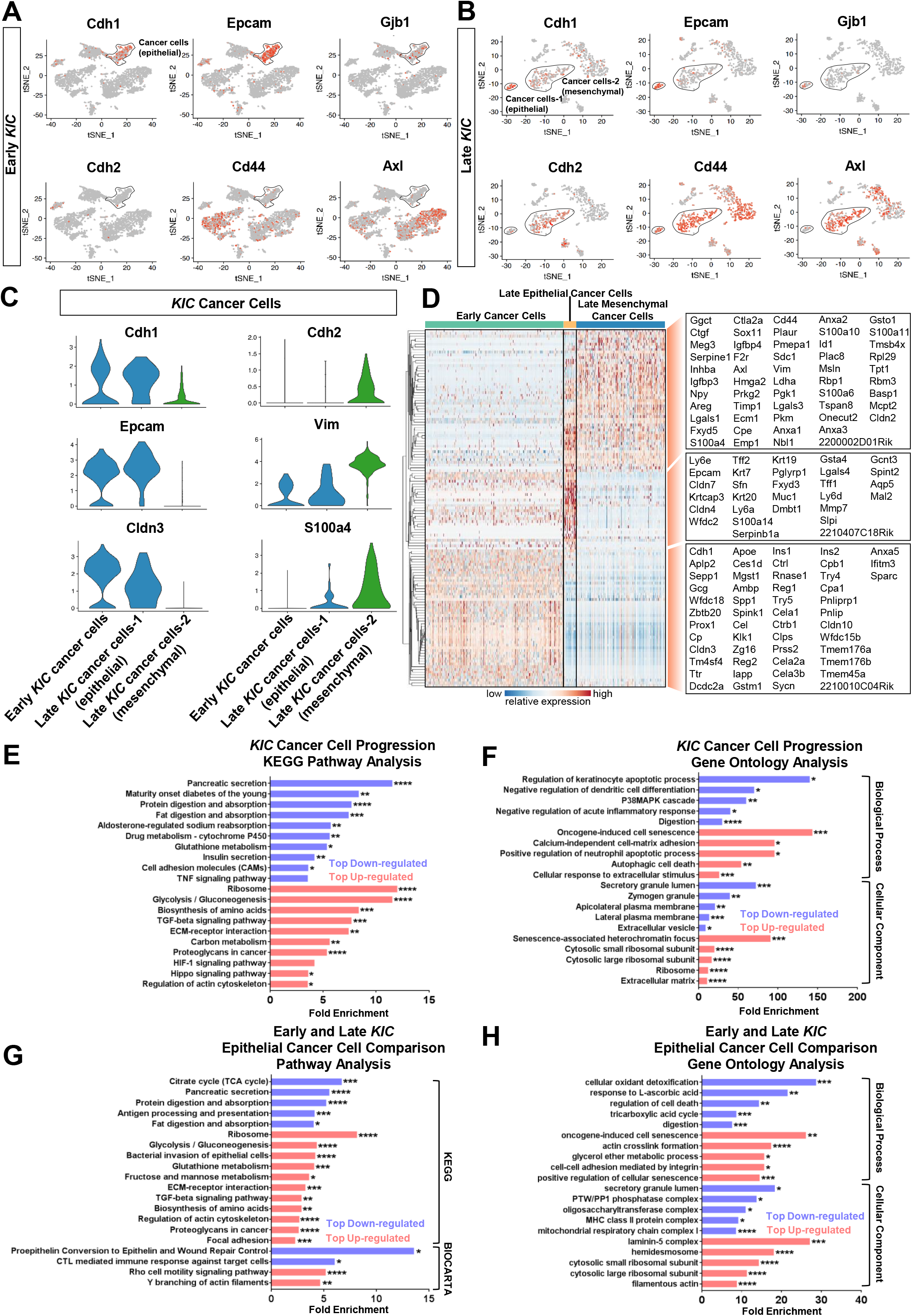
Analysis of early and late *KIC* cancer cell populations demonstrate the emergence of the mesenchymal cancer cell population as a late event. A) tSNE plots of the early *KIC* lesion demonstrated the expression of known epithelial markers in the sole cancer population (black outline). Mesenchymal markers were absent in this population. B) tSNE plots demonstrating the emergence of two cancer cell populations in the late *KIC* tumor. One cancer cell population expressed the epithelial markers (smaller population outlined in black) and a second expressed the mesenchymal markers (larger population outlined in black). C) Violin plots showing the high expression of epithelial markers in the early *KIC* cancer cell population and late *KIC* epithelial cancer cell population but not in the mesenchymal population. Mesenchymal markers were overexpressed in the mesenchymal cancer cell population but not in the early *KIC* or late *KIC* epithelial cancer cell populations. D) Single-cell profiling heatmap of all early and late *KIC* cancer cells displaying differentially expressed genes between the three cell populations. Gene names are listed in the boxes on the far right of the heatmap. Each column represents an individual cell and each row is the gene expression value for a single gene. E) KEGG pathway analysis and F) gene ontology analysis comparing all early *KIC* cancer cells against all late *KIC* cancer cells. Red bars are increased categories and blue bars are decreased categories. G) KEGG and BIOCARTA pathway analysis and H) gene ontology analysis comparing all early *KIC* cancer cells against only the late *KIC* epithelial cells. Red bars are categories increased in the late *KIC* and blue bars are decreased in the late *KIC*. (\*\*\*\**P* < 0.0001, \*\*\**P* < 0.001, \*\**P* < 0.01, \**P* < 0.05).

The hierarchical clustering of the top significant genes in each of the three cancer cell populations (epithelial cancer cells in early *KIC*, epithelial and mesenchymal cancer cell populations in late *KIC*) was performed (Fig. 2D). In addition, gene clusters from the cancer cell populations were subjected to pathway and gene ontology (GO) analysis. First, we compared cancer cells of the early *KIC* population to the total cancer cells of the late *KIC* and found that the most downregulated genes in late *KIC* cancer cells were associated with normal pancreatic function such as pancreatic secretion, digestion and absorption, and insulin secretion (Fig. 2E and F). Moreover, normal pancreatic acinar genes such as *Try4, Try5, Cela2a, Cela3b, Reg2*, and *Rnase1* were expressed at higher levels in early *KIC* cancer cells, while late *KIC* cancer cells expressed a higher level of the pancreatic ductal gene *Muc1* (Fig. 2D). This is suggestive of an ongoing acinar-to-ductal metaplasia (ADM) during tumor progression in this GEMM. In contrast, the most upregulated genes in late *KIC* cancer cells were associated with ribosome, glycolysis/gluconeogenesis, and amino acid biosynthesis, which is highly suggestive of increased translation and metabolically active cancer cells in established *KIC* tumors. Interestingly, pathways previously reported to be closely associated with the stroma and progression of PDA were also highlighted, such as ECM-receptor interaction [12], TGFβ [13], and hippo signaling pathways [14]. We then compared early *KIC* cancer cells with the late *KIC* epithelial cancer cell population to understand the mechanisms that promoted the progression of PDA in the epithelial cancer cell compartment. Interestingly, similar cell functions/signaling pathways were identified by comparing the two epithelial cancer cell populations (Fig. 2G and H). Taken together, these analyses objectively demonstrate an ADM state during the progression of *KIC* tumors and suggest that stroma-cancer cell interaction promotes the progression of PDA and cancer cell heterogeneity.

### Mesenchymal cancer cells exist in advanced PDA GEMMs with different diverse mutations

In addition to *KRAS* mutations, additional driver events are required for PDA progression [8], with *TP53* and *INK4A* being the second- and third-most commonly mutated genes in human PDA, respectively. As such, we sought to understand the effect of different secondary driver mutations on the phenotypes and heterogeneity of cancer cells. We performed scRNA-seq in another PDA GEMM, *KPfC* (*Kras^LSL-G12D^; Trp53^Flox/Flox^*; *Pdx1*^*Cre*/+^) (Fig. 3A). Consistent with late *KIC* tumors, two distinct cancer cell populations expressing *Krt18* and *Sox9* were noted in late *KPfC* (60-day-old) tumors (Fig. 3A and B), one marked by epithelial markers such as *Gjb1, Tjb1, Ocln*, and *Cldn3*, while the other was marked by mesenchymal markers such as *Vim, Cd44, Axl, S100a4*, and *Fbln2* (Fig. 3C and D). Epithelial and mesenchymal cancer cell populations in *KPfC* mice shared many genes in common with the corresponding populations in *KIC*; however, they also expressed unique gene signatures (Fig. 3F).

**Figure 3.**
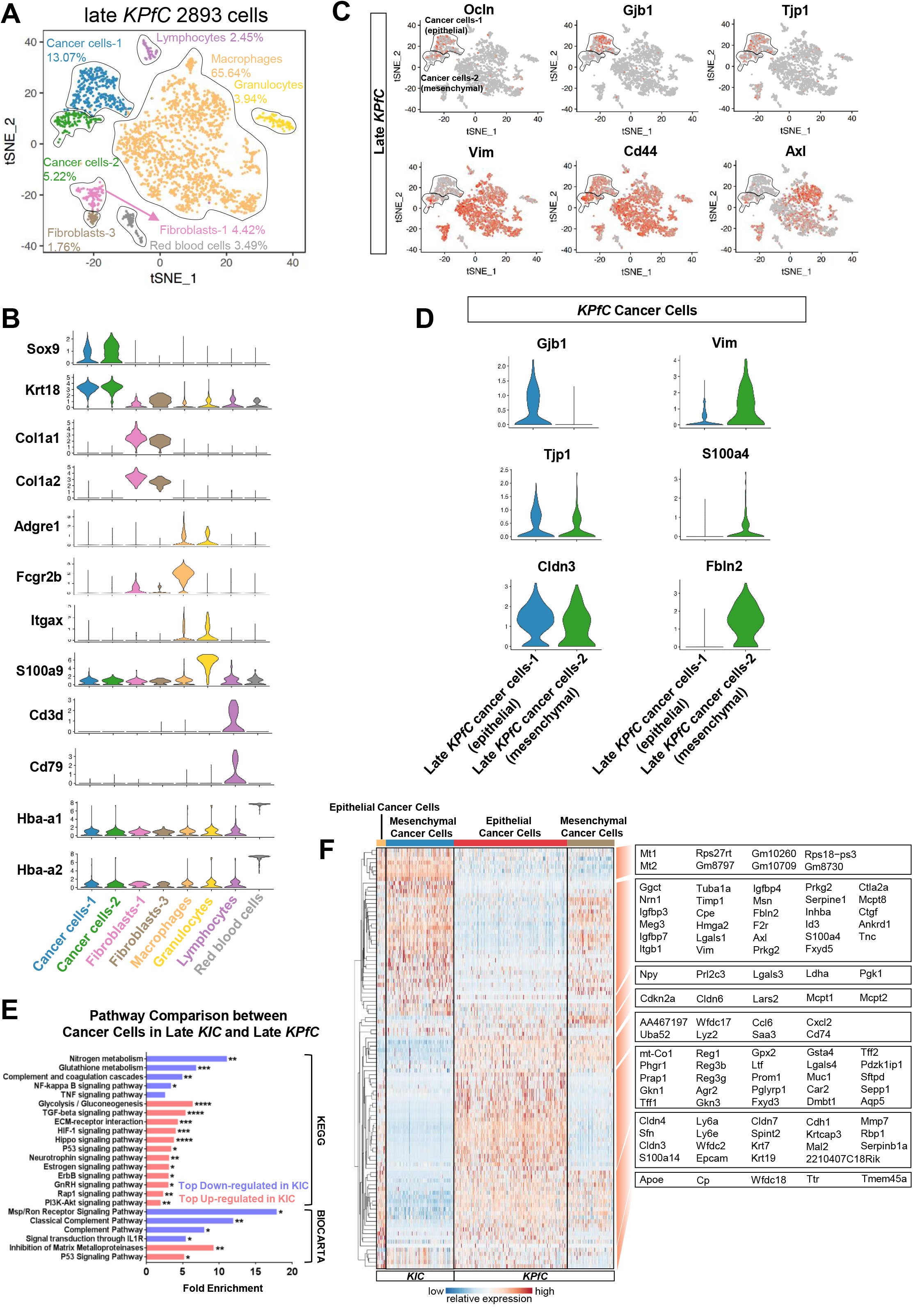
Comparison between cancer cells of *KIC* and *KPfC* tumors. A) tSNE plot of the late *KPfC* lesion displaying 2893 cells and 8 distinct cell populations. B) Stacked violin plots showing representative marker gene expression for each of the cell populations seen in the late *KPfC* lesion. C) Single-gene tSNE plots of the *KPfC* tumor displaying the presence of epithelial markers (*Ocln, Gjb1*, and *Tjp1*) in the epithelial cancer cell population (upper black outlined population) and mesenchymal markers in the mesenchymal cancer cell population (lower black outlined population). D) Violin plots showing the overexpression of epithelial markers in the epithelial cancer cell population and mesenchymal markers in the mesenchymal cancer cell population. E) KEGG and BIOCARTA pathway analysis comparing all late *KIC* to all late *KPfC* cancer cells. Red bars are categories increased in the late *KIC* and blue bars are increased in *KPfC*. F) Single-cell profiling heatmap comparing all cancer cells in the *KIC* versus all cancer cells in the *KPfC*. Each column represents an individual cell and each row is the gene expression value for a single gene. (\*\*\*\**P* < 0.0001, \*\*\**P* < 0.001, \*\**P* < 0.01, \**P* < 0.05).

We then compared the total cancer cell gene signatures between late *KIC* and late *KPfC* mice by KEGG and Biocarta pathway analysis methods, in an attempt to identify potential differences in cancer cell signaling pathways caused by the different secondary driver mutations. As expected, the p53 signaling pathway was upregulated in the *KIC* model by comparison to the *KPfC* model (Fig. 3E). The analyses of late *KIC* and late *KPfC* mice suggests that cancer cell heterogeneity is a late-stage tumor event that occurs in the setting of multiple secondary driver mutations. However, under the same oncogenic *Kras* mutation, different secondary driver mutations can potentially lead to different signaling pathways that drive PDA progression.

### Macrophage heterogeneity during PDA progression

We found a marked increase in the size of the macrophage population as PDA progressed from normal pancreas to early *KIC* and eventually late *KIC* tumors (Fig. 1B-D). We further characterized the macrophage compartment during PDA progression by subclustering macrophages in early and late *KIC* tumors, which revealed three transcriptionally distinct macrophage clusters in early *KIC* and two in late *KIC* (Fig. 4A and C).

**Figure 4.**
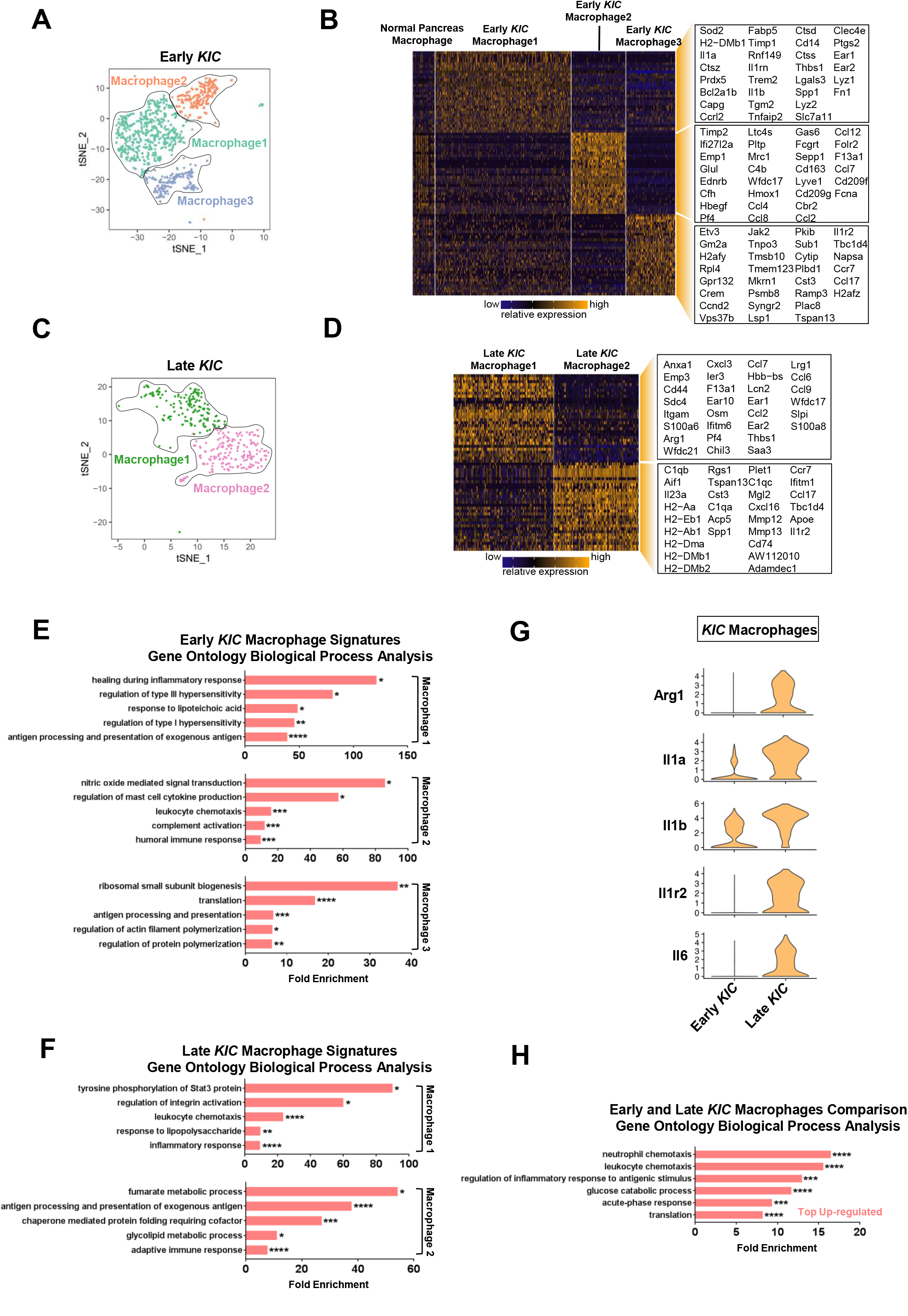
scRNA-seq analysis of *KIC* tumor progression reveals multiple subpopulations of macrophages. A) tSNE plot of three macrophage subpopulations in the early *KIC* tumor. B) Heatmap depicting the 30 top significantly overexpressed genes in each of the three early *KIC* macrophage subpopulations. Macrophages from the normal pancreas are displayed (far left group). Each column represents an individual cell and each row is the gene expression value for a single gene. C) tSNE plot representation of two macrophage subpopulations in the late *KIC*. D) Heatmap depicting the 30 top significantly overexpressed genes in each of the two late *KIC* macrophage subpopulations. Each column represents an individual cell and each row is the gene expression value for a single gene. E) GO analysis of biological processes in the three macrophage subpopulations seen in the early *KIC*. F) GO analysis of biological processes in the two macrophage subpopulations of the late *KIC*. G) Violin plots of the expression of inflammatory genes and *Arg1* comparing the macrophages in early and late *KIC*. H) GO analysis of biological processes that are upregulated in the late *KIC* macrophages relative to early *KIC* macrophages. (\*\*\*\**P* < 0.0001, \*\*\**P* < 0.001, \*\**P* < 0.01, \**P* < 0.05).

Macrophage population 1 in early *KIC* tumors was characterized by the expression of *Fn1, Lyz1, Lyz2, Ear1*, and *Ear2* as well as *Cd14* (Fig. 4B). Moreover, these macrophages specifically expressed high levels of the IL1 receptor ligands: *Il1a, Il1b*, and *Il1rn*. GO analysis suggested that this macrophage population was involved in healing during inflammation, the regulation of type I and III hypersensitivities, and antigen processing and presentation (Fig. 4E). In contrast, macrophage population 2 was noted to express an abundance of chemokines, including *Ccl2, Ccl4, Ccl7, Ccl8*, and *Ccl12*, as well as many complement-associated genes (Fig. 4B). Indeed, leukocyte activation, complement activation, and humoral response genes were the most significantly enriched GO categories in this macrophage population (Fig. 4E). The third macrophage population expressed *Ccl17* and *Ccr7* and was enriched in ribosomal small-unit biogenesis, translation, and antigen-processing functions (Fig. 4B and E). Importantly, macrophages in normal mouse pancreas weakly expressed genes found in macrophage population 2 and 3 from early *KIC* mice, suggesting that the normal pancreas macrophages could be noncommitted macrophages residing in tissue in the normal organ that are induced to adopt a distinct phenotype upon tumor initiation (Fig. 4B).

The late *KIC* tumor featured two macrophage subpopulations (Fig. 4C). Macrophage population 1 highly expressed genes such as *S100a8* and *Saa3*, which have been shown to be expressed in lipopolysaccharide-treated monocytes [15]. Moreover, numerous chemokines were elevated in this population such as *Ccl2, Ccl7, Ccl9, Ccl6, Cxcl3*, and *Pf4* (Fig. 4D). GO analysis revealed this population is likely associated with Stat3 activation, leukocyte chemotaxis, and response to lipopolysaccharide and inflammatory stimuli (Fig. 4F). These data suggest that macrophage population 1 was inflammatory in nature. Macrophage population 2 of late *KIC* tumors was rich in MHC-II antigen presentation molecules: *Cd74, H2-Aa, H1-Ab1, H2-Dma, H2-Dmb1, H2-Dmb2, and H2-Eb1* (Fig. 4D), and GO analysis highlighted antigen presentation and adaptive immune response pathways as being elevated (Fig. 4F). Consistently, in late *KPfC* tumors, we also observed two distinct populations of macrophages with similar features (Supplementary Fig. 3). Interestingly, we did not observe a macrophage population in late tumors that correlated with macrophage population 1 from the early tumors, suggesting that this population might undergo negative selection or a differentiation into inflammatory and/or MHC-II–rich macrophages during tumor progression.

We also compared the features of the total macrophage clusters between early and late *KIC* tumors and observed a substantially enhanced macrophage inflammatory signature as the tumor progressed (Fig. 4G). A wide variety of inflammatory genes increased, including *Il1a, Il1b, Il1r2*, and *Il6*. GO analysis of this gene list highlighted leukocyte chemotaxis and inflammatory response functions as increased in advanced *KIC* tumors (Fig. 4H). These data suggest that PDA progression is characterized by an increase in inflammatory features in macrophages.

### Fibroblast heterogeneity during PDA progression

In normal pancreas and early *KIC* tumors, we had identified three distinct populations of fibroblasts, while in late *KIC* only two fibroblast populations were noted (Fig. 1B-D). To ascertain the relationship between these fibroblast populations and the dynamics of their phenotypic changes during PDA progression, we projected fibroblasts from the three analyses onto a single tSNE plot and applied a graph-based clustering algorithm (Fig. 5A) which revealed three distinct molecular subtypes of fibroblasts in the normal pancreas, early *KIC* tumors, and late *KIC* tumors. The overlay demonstrates that the normal pancreas and early *KIC* tumors contained all three fibroblast subtypes while the late *KIC* contained only two (Fig. 5A), confirming our initial analysis (Fig. 1B-D). Specifically, this analysis demonstrated that fibroblast population 1 (FB1) and fibroblast population 3 (FB3) found in normal and early *KIC* pancreas were present in the late *KIC* tumor whereas fibroblast population 2 (FB2) was absent.

**Figure 5.**
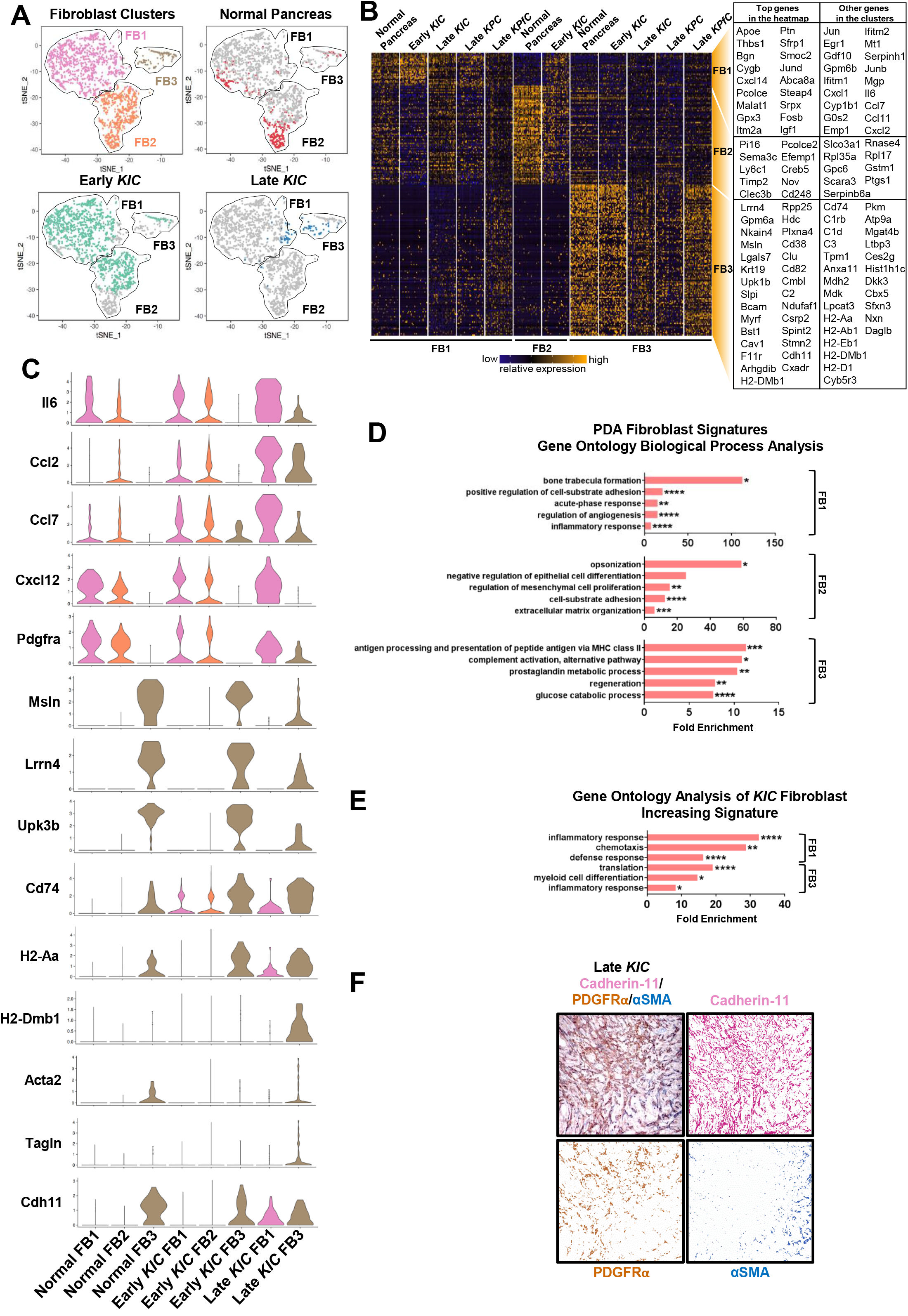
Analysis of fibroblasts during PDA progression reveals multiple molecular subtypes. A) All fibroblasts from the normal pancreas, early *KIC* tumors, and late *KIC* tumors were projected onto a single tSNE plot with the FB1, FB2, and FB3 populations distinguished by pink, orange, and brown, respectively (upper left panel). Normal pancreas fibroblasts were highlighted in red (upper right panel), early *KIC* fibroblasts in green (lower left panel) and late *KIC* fibroblasts in blue (lower right panel). Normal pancreas and early *KIC* contained fibroblasts in all three groups whereas the late *KIC* had only FB1 and FB3. B) Heatmap displaying the top significant genes (cutoff: *P* < 10^-40^) for each of the three fibroblast populations. Thirty random cells from each fibroblast population are displayed. All three late-cancer GEMMs (late *KIC*, *KPfC*, and *KPC*) display only FB1 and FB3 fibroblast populations. C) Violin plots demonstrating representative marker genes for each fibroblast subtype: FB1 overexpressed cytokines and *Pdgfra*. FB3 overexpressed mesothelial markers, myofibroblast markers, MHC-II molecules and *Cdh11*. D) Gene ontology analysis of the top biological processes in each of the three fibroblast subtypes. E) GO analysis of genes upregulated in late FB1 and FB3 compared to early FB1 and FB3 in *KIC*, respectively. F) Immunohistochemical analysis of PDGFRα, αSMA, and CDH11 stained serially on the same slide. Colors were deconvoluted into a single color layer. PDGFRα and αSMA staining were mutually exclusive, whereas CDH11 was a pan-CAF marker (magnification: 20x). (\*\*\*\**P* < 0.0001, \*\*\**P* < 0.001, \*\**P* < 0.01, \**P* < 0.05).

In the normal pancreas, FB1, FB2, and FB3 made up 35.4%, 56.9% and 7.7% of the total fibroblasts, respectively (Supplementary Fig.4A). In early *KIC* tumors, although the total fibroblasts expanded (Fig. 1C), the ratios of each fibroblast population remained similar. Furthermore, in the late *KIC* tumors, FB1 and FB3 were present in nearly equal proportions of 46.5% and 53.5%, respectively (Supplementary Fig.4A). Each fibroblast population was characterized by distinct marker genes. For example, FB1 markedly expressed *Cxcl14, Ptn*, and several genes mediating insulin-like growth factor signaling such as *Igf1, Igfbp7*, and *Igfbp4*. FB2 specifically expressed *Nov*, a member of the CCN family of secreted matricellular proteins [16] as well as *Pi16*, which has been shown to be expressed in fibroblast populations in various tissue types [17], in addition to *Ly6a* and *Ly6c1*. FB3 showed distinct expression of mesothelial markers such as *Lrrn4, Gpm6a, Nkain4, Lgals7*, and *Msln* [18] in addition to other genes previously shown to be expressed in fibroblasts such as *Cav1, Cdh11*, and *Gas6* [19–21].

Hierarchical clustering of the most significant genes for each fibroblast subtype confirmed the persistence of FB1 and FB3 during the progression of PDA (Fig. 5B) and that they exist across different advanced-stage PDA GEMMs (*KPC* and *KPfC*), suggesting a consistent cell of origin. Interestingly, the gene expression heatmap also indicated that the FB2 population started to move toward an FB1-like expression profile in early *KIC* tumors, suggesting FB1 and FB2 might converge into a single CAF population with FB1 features by late invasive disease. Of note, *Il6, Ccl2, Ccl7, Cxcl12*, and *Pdgfra* were expressed in FB1 and FB2 in the normal pancreas and early *KIC* tumors, and showed greater expression in FB1 of late *KIC* (Fig. 5C). In contrast, the myofibroblast markers *Acta2* and *Tagln* were expressed by a portion of FB3. These data support the presence of previously described, mutually exclusive, inflammatory (FB1) and myofibroblastic (FB3) CAF subtypes [22–24]. Interestingly, FB3 also expressed numerous MHC-II–associated genes (Fig. 5C). GO analysis suggested that FB1 was involved in an acute phase response and inflammatory response, FB2 was more associated with physiological functions of fibroblasts, while FB3 had antigen processing and presentation through the MHC-II pathway and had complement activation functions (Fig. 5D). Furthermore, we analyzed genes that increased in FB1 and FB3 during PDA progression, and found that FB1 showed a progressive increase in the expression of genes associated with inflammatory response and chemotaxis (Fig. 5E and Supplementary Fig. 4B) while FB3 genes displayed increased function on translation during disease progression, possibly due to enhanced antigen processing activity. These data suggest that FB1 is an inflammatory population and the inflammatory feature increases during PDA progression, while FB3 consists of the well-studied myofibroblast population, and displays an enrichment for class 2 MHC genes.

We also found that some genes essentially exclusive to FB3 in the normal and early *KIC* pancreas became expressed in FB1 and FB3 populations in late *KIC*, marking these genes as potential global fibroblast markers in advanced PDA. One such gene was *Cdh11* (Fig. 5C). We validated these data by immunohistochemistry. We found in late *KIC* tumors, stromal staining for αSMA and PDGFRα were nearly mutually exclusive, whereas CDH11 showed uniform staining across all morphologically discernable fibroblasts (Fig. 5F). Taken together, these data provide the first *in vivo* description of all CAF populations during PDA progression.

### Mesenchymal cancer cells and CAFs show evidence of increased epigenetic regulation and super-enhancer activity in advanced PDA

Unique molecular identifiers (UMI) serve to barcode each input mRNA molecule during cDNA library generation, enabling the determination of initial transcript number even after cDNA library amplification [25]. We compared UMI counts across all cell types between early and late *KIC* tumors (Fig. 6A and B). In early lesions, there was a marked increase in UMI in the beta islet cells (median: 2849, range: 1322-12,857), which might indicate that increased transcriptional activity is a means by which the endocrine requirements of these cells are met. No other cell population in the early *KIC* tumor displayed this level of UMI. The early *KIC* cancer cells displayed a relatively low UMI count (median: 1979, range: 1163-7735). In contrast, the mesenchymal cancer cell population in the late *KIC* tumor displayed a marked increase in total UMI count with a median count of 18,334 and range of 4433-50,061 (Fig. 6C). The epithelial cancer cells in the late *KIC* also displayed an increased UMI, albeit to a far lesser degree than the mesenchymal cancer cell population (median: 10,368, range: 4940-30,440).

**Figure 6.**
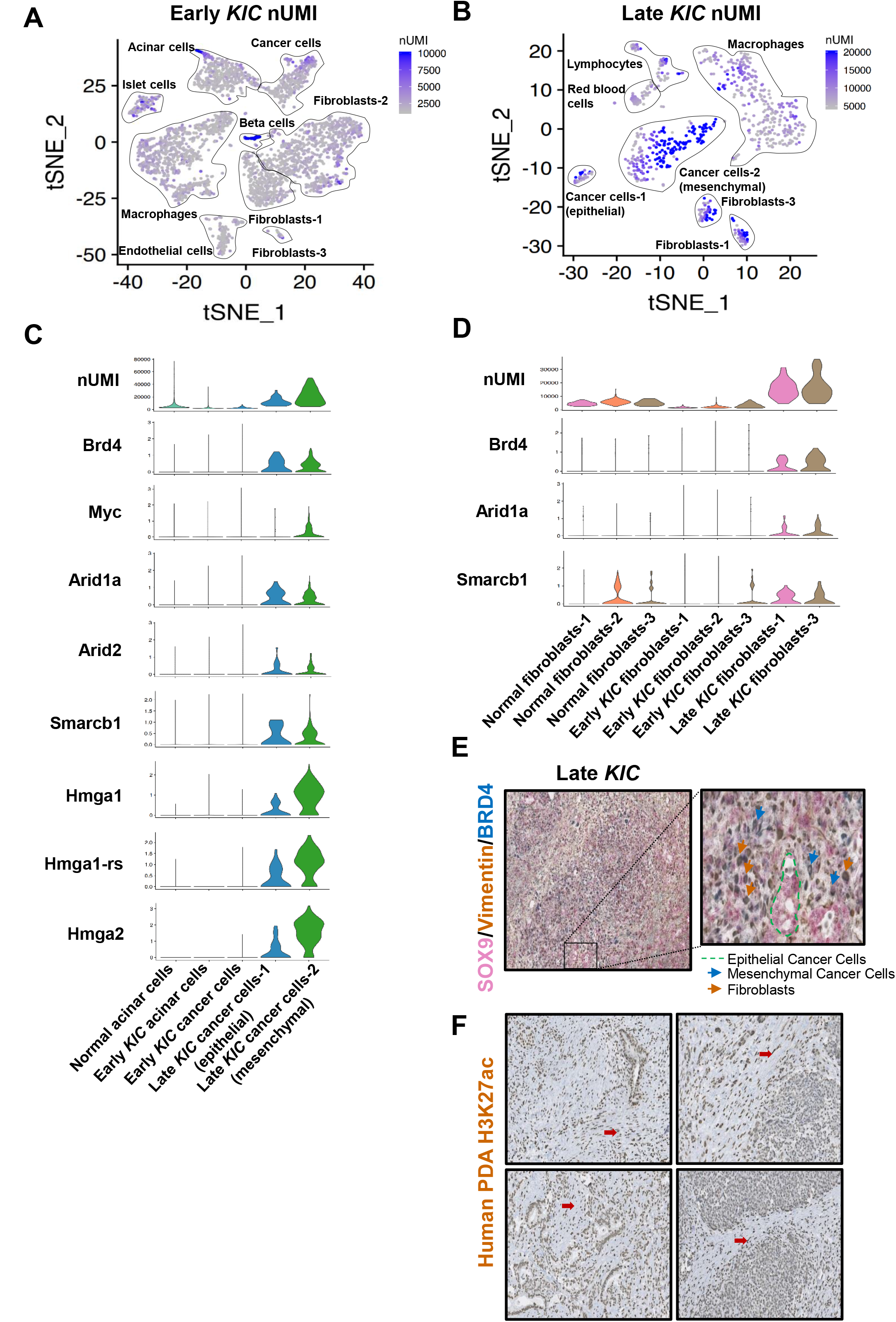
Analysis of transcriptional activity in different stages of PDA reveals differential epigenetic and super-enhancer activity in distinct tissue compartments. A) tSNE plot of UMI counts in the early and B) late *KIC*. C) Violin plots of epigenetic regulatory genes in the cancer cell populations of normal pancreas, early *KIC*, and late *KIC*. D) Violin plots of epigenetic regulator genes in the normal, early fibroblast, and late fibroblast populations showing their upregulation in CAFs. E) Sequential triple immunohistochemical staining on the same late *KIC* tumor section for cancer cells (SOX9, pink), mesenchymal cells (vimentin, brown) and super-enhancer activity (BRD4, blue). Well-differentiated ductal epithelium stained solely for SOX9 (green outline). Mesenchymal cancer cells (blue arrows) and CAFs (brown arrows) both show co-staining with BRD4. F) Immunohistochemical analysis of human PDA whole tissue sections using the H3K27ac antibody. These representative figures from four different human PDAs demonstrate the 3+/3+ staining in the stromal fibroblasts (red arrows) with 1-2+ staining in the cancer epithelium (magnification: 20x).

We reasoned that the increased transcriptional activity may be associated with increased activity of epigenetic regulation as well as super-enhancer [26]. BRD4 belongs to the bromodomain family of transcriptional regulators and is a key regulator of super-enhancer activity [27]. Prior studies have shown that MYC activity is promoted by super-enhancer activity in PDA [28]. We found that in late *KIC* and *KPfC* tumors, *Brd4* was expressed highly in epithelial and mesenchymal cancer cells while *Myc* was expressed mainly in the mesenchymal cancer cell population (Fig. 6C, Supplementary Fig. 6). In addition, several genes encoding high-mobility group A proteins (*Hmga1, Hmga1-rs, Hmga2*) were markedly expressed in late *KIC* and *KPfC* mesenchymal cancer cells. HMGA proteins are chromatin-associated proteins that regulate transcriptional activity, including enhancesome formation [29]. Lastly, critical components of the SWI/SNF complex (*Smarcb1, Arid1a, Arid2*), which are essential in nucleosome remodeling and transcriptional regulation [30], were also expressed highly in epithelial and mesenchymal cancer cells of the late *KIC*, but not cancer cells in the early *KIC* lesion. Taken together, these data provide multiple lines of evidence to suggest that the transcript load of a more aggressive mesenchymal cancer cell population is increased relative to cancer cells in early lesions or epithelial cancer cells in advanced PDA.

Interestingly, we also noted that fibroblasts in late *KIC* tumors also showed an increased UMI (median: 14,538, range: 4461-37,497). They also displayed an increased expression of super-enhancer and other epigenetic transcriptional regulator genes in contrast to fibroblasts from normal mouse pancreas or early *KIC* pancreas (Fig. 6D). These data are suggestive of increased super-enhancer and transcriptional activity as normal pancreas fibroblasts become CAFs.

We validated these single-cell RNA expression data using three-color immunohistochemical analysis of late *KIC* tumors: SOX9 was used as a pan-cancer cell marker, vimentin as a mesenchymal marker, and BRD4 was a surrogate marker for super-enhancer activity. We identified positive co-staining for vimentin and Brd4 in CAFs, positive triple-staining (vimentin+/Sox9+/Brd4+) in mesenchymal cancer cells, and single staining of Sox9 in epithelial cancer cells that localized to more differentiated, duct-like structures in the advanced tumors (Fig. 6E). Next, we performed immunohistochemical analysis on 16 whole tumor human pancreatic cancer sections using an antibody against H3K27ac, a commonly accepted marker of increased gene regulatory element activity [26, 31]. The malignant epithelium and stromal fibroblasts were scored separately. These analyses showed markedly positive 3+/3+ staining in the stromal fibroblasts of all whole tumor sections (Fig. 6F). In 6/16 cancer epithelia the score was 1+ and 10/16 scored 2+, with no samples showing a cancer epithelial scoring of 3+. Taken together, these are the first data indicating differential super-enhancer activity in distinct tissue compartments of PDA.

## Discussion

We have carried out an scRNA-seq of different stages of the *KIC* GEMM, in addition to late *KPfC* and *KPC* tumors in an effort to agnostically profile the phenotypic changes of cancer and stromal cells during PDA progression. We have established the emergence of a mesenchymal cancer cell population as a late-stage tumor event and have identified novel features of different macrophage and fibroblast populations. This significantly improves our understanding of PDA progression and lays the foundation for the development of novel therapeutic approaches.

PDA pathogenesis involves metaplasia of normal acinar cells to ductal epithelium, which in turn undergo neoplastic transformation in a KRAS-driven manner [32]. Malignant ductal epithelium may then assume more aggressive, mesenchymal features as the disease progresses. In this study, mesenchymal cancer cell populations were noted in late-stage tumors. Our data support a model in which mesenchymal features of cancer cells are acquired later in the disease process, although others have argued that this can be one of the earliest events in PDA [33]. Mesenchymal cancer cell populations have been studied extensively in pancreatic cancer mouse models and has been shown to be critical to chemotherapeutic resistance while their contribution to metastasis has been more controversial [34, 35]. Mesenchymal cancer cells have previously demonstrated an increased protein anabolism and activation of the endoplasmic reticulum–stress-induced survival pathways in a PDA GEMM [36].

Indeed, in the late *KIC* model, ribosomal pathways were the most significantly upregulated pathways in cancer cells (Fig. 2E-H). It is likely that the demand for increased ribosomal activity stems from high transcriptional activity governed by epigenetic mechanisms in the mesenchymal cancer cells, as we also saw markedly increased UMI counts in this population (Fig. 6B). The bromodomain and extraterminal (BET) family of proteins such as BRD4, which is markedly upregulated in cancer cells and fibroblasts of late-stage PDA (Fig. 6C and D), serve to recruit regulatory complexes to acetylated histones at enhancer sites, resulting in increased transcription [37]. Previously, a combination approach using a BET protein inhibitor and a histone deacetylase inhibitor led to near-complete tumor regression and improved animal survival in a PDA GEMM [28]. Super-enhancer activation has recently been shown to be fundamental in the pathophysiology of a variety of neoplasms [38] and is intimately associated with *Hmg2a* in PDA, as super-enhancer attenuation has been demonstrated to downregulate *Hmg2a* expression and the growth of PDA cells in a three-dimensional *in vitro* model [39]. Our study is the first to demonstrate that these epigenetic regulatory mechanisms in PDA are present in specific tissue compartments, namely mesenchymal cancer cells and CAFs. Future efforts to target super-enhancer activity in PDA should consider distinct tissue compartments governing the sensitivity and resistance to novel therapeutics.

Our data revealed two molecular subtypes of macrophages in advanced PDA (Fig. 4). One expressed numerous chemokine and inflammation-associated genes while the other was rich in MHC-II–associated genes. In a previous study, MHC-II–positive macrophages were isolated from orthotopic breast tumors and highly expressed CCL17, consistent with our data [40]. In parallel with our study, MHC-II^low^ macrophages were found to be highly enriched for numerous chemokines. Moreover, in an orthotopic hepatoma mouse model, an early MHC-II+ macrophage population appeared to suppress tumor growth but an MHC-II^low^ macrophage population became the predominant macrophage population as the tumor progressed, resulting in a protumor phenotype [41]. Nonetheless, to confirm their pathophysiological significance, functional studies are required in which inducible selective ablation [42] is performed on the two late-stage PDA macrophage subpopulations using specific markers we have identified in this study. Zhu and colleagues [43] have shown that bone marrow-derived monocytes make up approximately 80% of MHC-II–positive macrophages in a PDA GEMM whereas MHC-II–negative macrophages in normal pancreas and PDA were shown to be maintained independently of monocyte contributions. Monocyte-independent MHC-II^low^ tissue resident macrophages expanded during tumor progression and contributed to PDA growth and survival. Conversely, Sanford and colleagues [44] have shown that monocytes can give rise to a pro-inflammatory macrophage population in a PDA mouse model, which when antagonized with neutralizing antibodies against CCR2, resulted in decreased tumor growth and reduced metastases *in vivo* [44]. These data highlight the need for an scRNA-seq study on macrophage populations in PDA GEMMs with labelled bone marrow replacement to reconcile these discrepancies.

More importantly, in the studies of tumor-associated macrophages, inflammatory chemokines are commonly used to indicate an M1 type of macrophage, which are normally associated with immune-stimulatory functions. Nevertheless, our study indicates that a distinct M1/M2 macrophage phenotype is not readily discernable at the single-cell level. Instead, as PDA progresses, an inflammatory feature is substantially increased, and this accompanies an increase of an important M2 macrophage marker, ARG1 (Fig. 4F and H). This raises questions on the M1/M2 classification system, as the inflammatory feature is associated with the progression of PDA. Future studies should focus on the function of these inflammatory macrophages in PDA in addition to validating markers for macrophage classification.

While numerous studies have generally shown that CAFs are tumor-promoting in the biology of PDA and other carcinomas [45, 46], recent studies have found that the function of CAFs in PDA biology are more varied. Özdemir and colleagues [42] demonstrated that the depletion of αSMA+ cells from the microenvironment in a PDA GEMM resulted in shortened survival and poorly differentiated tumors [42], and low myofibroblast tumor content was shown to be associated with worse survival in human PDA sections. These data prompted a paradigm shift whereby certain CAFs may function to constrain, rather than promote, PDA. Moreover, until recently, the molecular heterogeneity of CAFs in PDA has not been well-appreciated. The primary attempt to characterize fibroblast heterogeneity in PDA demonstrated that mouse pancreatic stellate cells (PSCs) could be induced to express αSMA *in vitro* when directly co-cultured with primary mouse PDA cells in an organoid co-culture system [22]. These myofibroblastic CAFs were designated as “myCAFs.” This was distinct from IL6+ fibroblasts that were produced *in vitro* when PSCs were indirectly co-cultured with mouse PDA organoids through a semi-permeable membrane. The IL6+ fibroblasts were also positive for PDGFRα and numerous other cytokines and therefore termed inflammatory CAFs or “iCAFs.” The immunohistochemistry of human and mouse PDA tissue showed distal IL6+ stroma as a distinct population from the peritumoral αSMA+ stroma. Subsequent studies in PDA GEMMs demonstrated that the iCAF population can mediate pro-tumorigenic properties and is a potential therapeutic target in an attempt to sensitize PDA to immunotherapeutic strategies [23, 24].

Our current study is the first to demonstrate the existence of three distinct molecular subtypes of fibroblasts in the normal mouse pancreas, which in turn gave rise to two distinct subtypes of CAFs that were largely conserved across three different PDA GEMMs. We noted that FB1 expressed insulin-like growth factor signaling genes (*Igfbp7, Igfbp4*, and *Igf1*) in addition to *Pdgfra, Cxcl12, Il6*, and several other cytokines (*Ccl11, Ccl7, Ccl2*, and *Csf1*). We propose that our FB1 population is the previously described iCAF population and hence likely pro-tumorigenic. Conversely, the FB3 population was positive for the myofibroblast markers *Acta2* and *Tagln*, and therefore most closely represents the previously described myCAF population. Importantly, our agnostic approach did not identify any further putative CAF populations and so we support the two-CAF model proposed by Öhlund and colleagues [22].

In summary, this report systematically outlines the cellular landscape during the progression of PDA and highlights the cellular heterogeneity in PDA pathogenesis. As such, future targeted therapeutic strategies should be developed with their intended target subpopulation in mind.

## Address requests for reprints to

Rolf A Brekken, 6000 Harry Hines Blvd., Dallas, TX 75390-8593.

## Acknowledgements

We thank the McDermott Center Next-Generation Sequencing Core at UT Southwestern for preparing and sequencing the single-cell RNA-seq libraries. We thank Dr Jeon Lee (UT Southwestern) for assistance in pre-processing of scRNA-seq data. We also thank Dave Primm of the UT Southwestern Department of Surgery for help in editing this article.

## Ethics approval

Mice were used in this study according to UT Southwestern institutional guidelines and approved by the institutional animal care and use committee at UT Southwestern Medical Center. All human samples were procured through the UT Southwestern tissue management shared resource and approved through the UT Southwestern institutional review board.

## Data sharing statement

There are no additional unpublished data from this study.

---

**Supplementary Figure 1.**
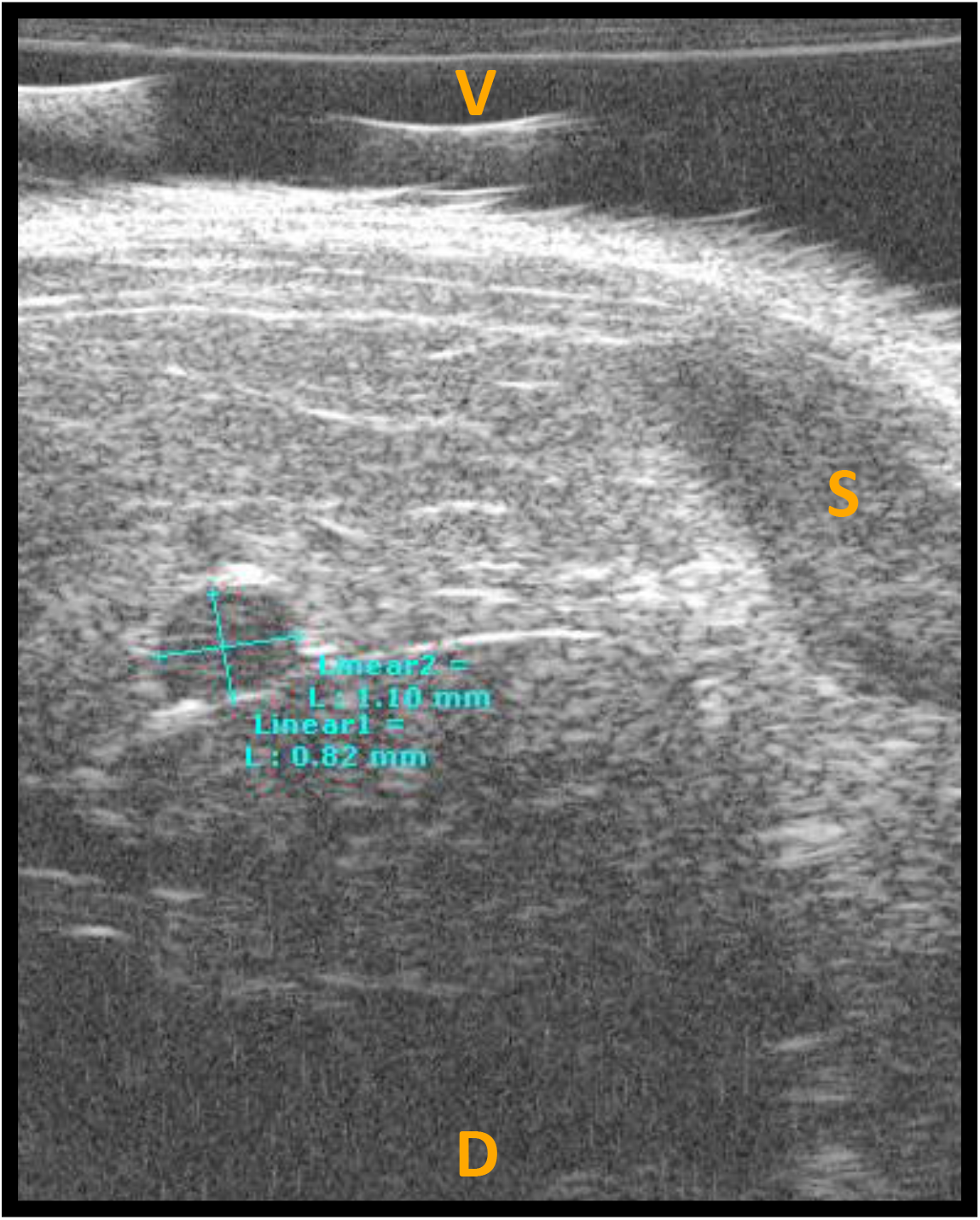
Ultrasound image of early *KIC* mouse (39 days old), 1 day prior to sacrifice. Two-dimensional measurement of the neoplastic lesion is denoted in teal (1.10 mm x 0.82 mm). V = ventral surface, D = dorsal surface, S = spleen.

**Supplementary Figure 2.**
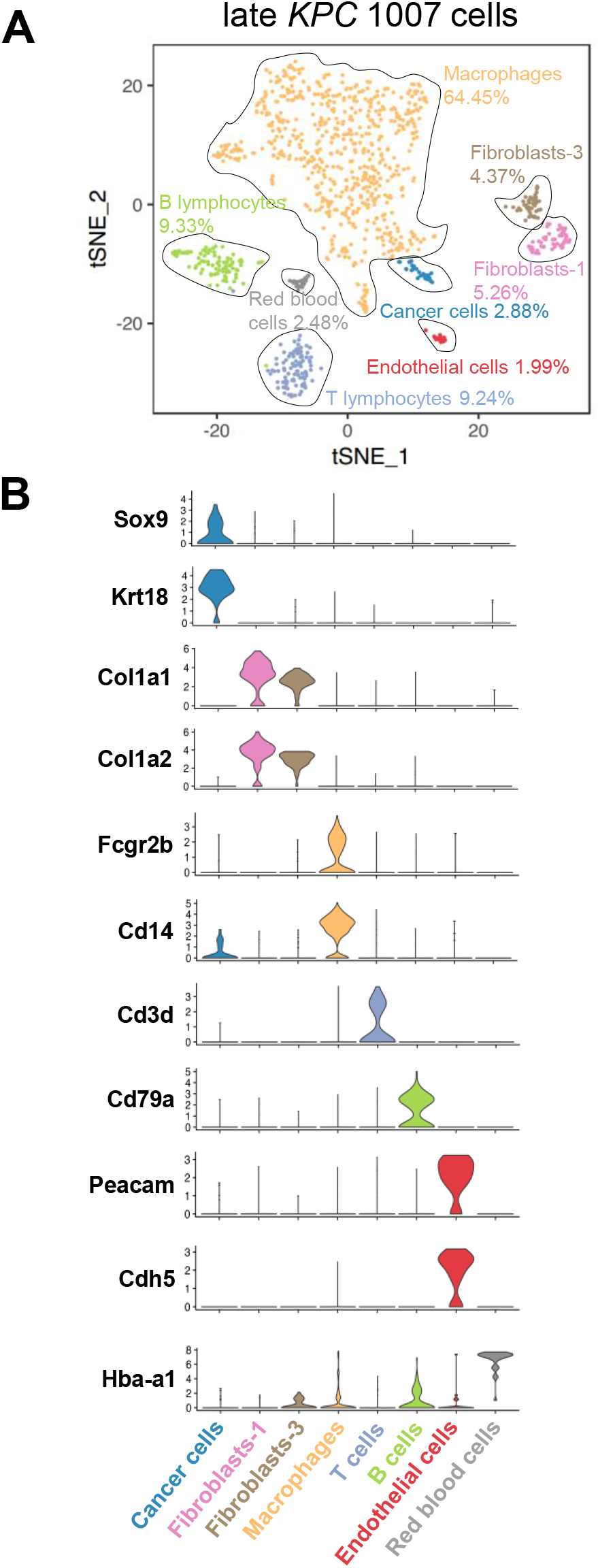
A) tSNE plot of *KPC* (6 months) displaying 1007 cells making up 8 distinct cell populations as indicated. B) Stacked violin plots of marker genes and UMI for each of the 8 *KPC* populations.

**Supplementary Figure 3.**
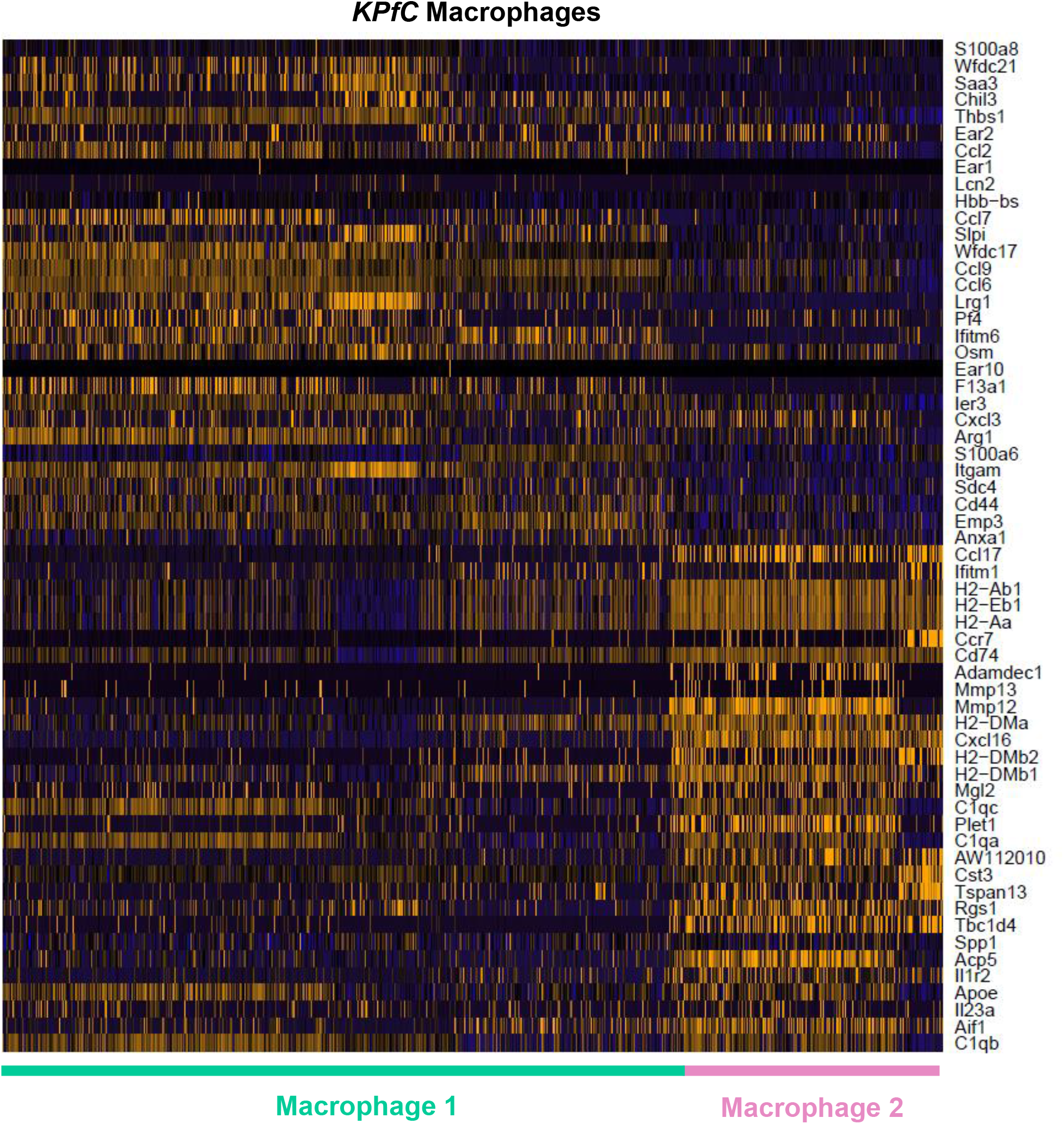
Heatmap depicting gene expression levels (horizontal) of single cells (vertical) in the macrophage populations of the *KPfC* GEMMs. Proinflammatory (Macrophage 1) and MHC-II-rich (Macrophage 2) subtypes are noted below the heatmap.

**Supplementary Figure 4.**
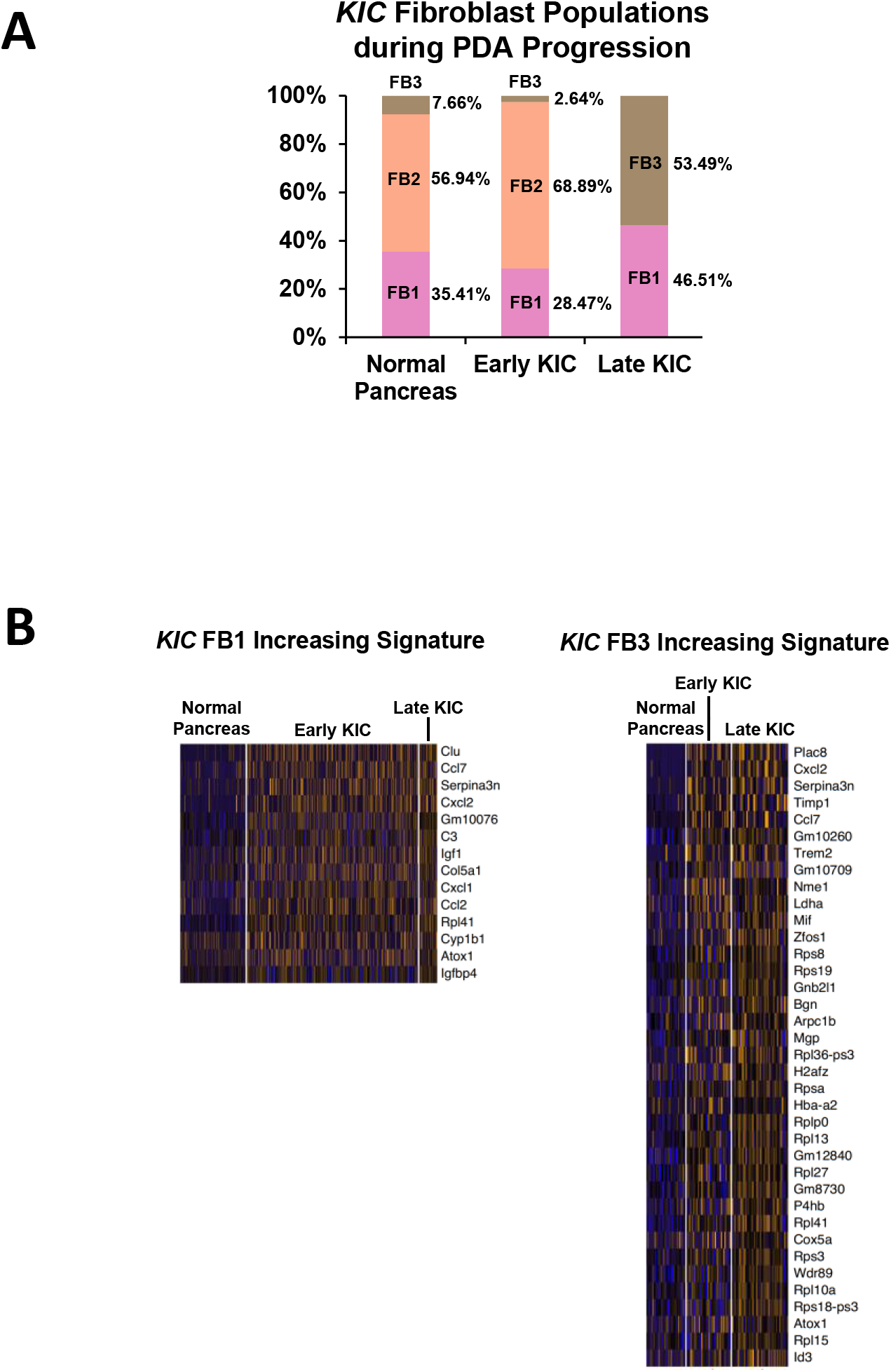
A) The relative proportions of FB1, FB2, and FB3 in the normal mouse pancreas, early *KIC* lesions, and late *KIC* lesions. B) Genes increased most significantly in the FB1 and FB3 populations as the normal pancreas progressed to early *KIC* and then to late *KIC*.

**Supplementary Figure 5.**
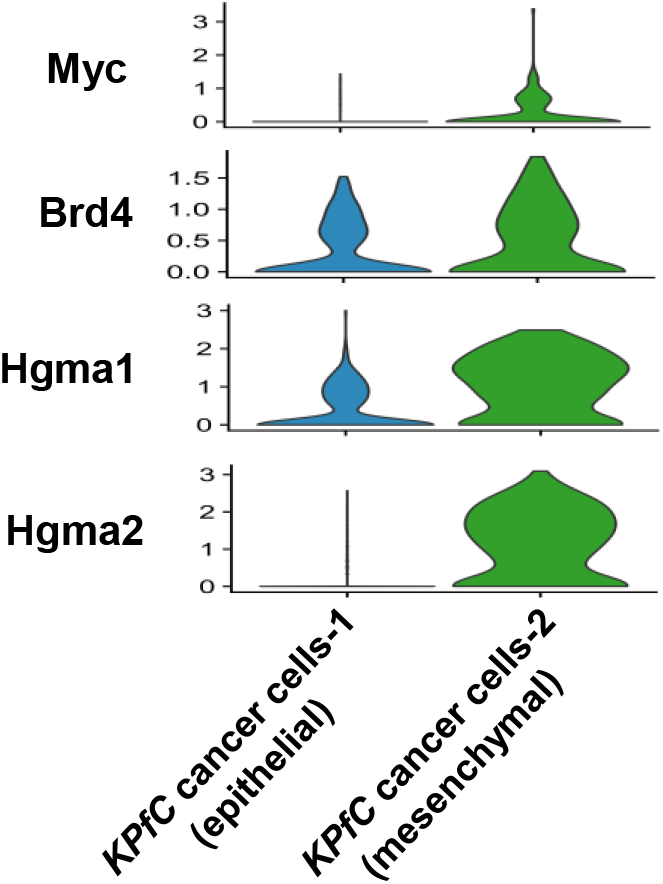
Violin plots depicting gene expression of *Myc* and epigenetic regulatory genes in the *KPfC* epithelial and mesenchymal cancer cell populations.

**Author contributions**
ANH: study concept and design, acquisition of data, analysis and interpretation of data, drafting of the manuscript. HH: study concept and design, acquisition of data, analysis and interpretation of data, drafting of the manuscript. ZW: bioinformatics analyses, interpretation of data. KP: pathologic interpretation and scoring of human tissue samples. WD: acquisition of data. JH: bioinformatics analyses, critical review of the manuscript. AM: bioinformatics analyses, interpretation of data, critical review of the manuscript. EO: bioinformatics analyses, interpretation of data, critical revision of the manuscript. UV: study concept and design, critical revision of the manuscript. RAB: study concept and design, interpretation of data, drafting of the manuscript.

## Abbreviations

ADM: acinar-to-ductal metaplasia
BET: bromodomain and extraterminal
CAF: cancer-associated fibroblast
FB: fibroblast population
GEM: gel bead in emulsion
GEMM: genetically engineered mouse model
GO: gene ontology
PDA: pancreatic ductal adenocarcinoma
PSC: pancreatic stellate cell
scRNA-seq: single-cell RNA sequencing
UMI: unique molecular identifiers

